# Inserting a Neuropixels probe into awake monkey cortex: two probes, two methods

**DOI:** 10.1101/2023.06.26.546631

**Authors:** Tomoyuki Namima, Erin Kempkes, Bob Smith, Anitha Pasupathy

**Affiliations:** Department of Biological Structure and Washington National Primate Research Center, University of Washington, Seattle, WA 98195, USA; Graduate School of Frontier Biosciences, Osaka University, and Center for Information and Neural Networks, National Institute of Information and Communications Technology, Suita, Osaka, 565-0871, Japan; Washington National Primate Research Center Instrumentation Services, University of Washington, Seattle, WA 98195, USA

**Keywords:** Nonhuman primate neurophysiology, high-density silicon probe, multi-contact linear probes, chronic recording chamber

## Abstract

Neuropixels probes have revolutionized neurophysiological studies in the rodent, but inserting these probes through the much thicker primate dura remains a challenge. Here we describe two methods we have developed for the insertion of two types of Neuropixels probes acutely into the awake monkey cortex. For the fine rodent probe, which is unable to pierce native primate dura, we developed a dural-eyelet method to insert the probe repeatedly without breakage. For the thicker NHP probe, we developed an artificial dura system to insert the probe. We have now conducted successful experiments in 3 animals across 7 recording chambers with the procedures described here and have achieved stable recordings over several months in each case. Here we describe our hardware, surgical preparation, methods for insertion and methods for removal of broken probe parts. We hope that our methods are of value to primate physiologists everywhere.

## 1. Introduction

The development and use of the Neuropixels probe with its high channel count and dense contact spacing has transformed electrophysiological methods, facilitating the study of hundreds of neurons at single neuron resolution (Jun et al., 2017; Steinmetz et al., 2018). In rodents, these probes have been used successfully either via acute insertion (Jun et al., 2017; Steinmetz et al., 2021) or chronic implantation (Juavinett et al., 2019; Luo et al., 2020; van Daal et al., 2021; Ghestem et al., 2023). In the awake macaque monkey however, their adoption has been limited. The original version of the probe developed for use in rodents (Neuropixels 1.0 aka. phase 3B2, IMEC) is fine, with a cross-section of 70 μm x 24 μm. This probe cannot pierce the thick macaque dura even when sharpened. The newer non-human primate (NHP) probes (Neuropixels NP1010, IMEC) on the other hand, are thicker (70 μm x 97 μm) and, when sharpened, can pierce primate dura soon after craniotomy. However, as the dura thickens over a 2-4 week period, the probe is unable to pierce the dura and a durotomy is required to continue recording from that area. Because both the rodent and NHP probes have a 10 mm shank length, neither can be inserted with traditional guide tube methods often employed with longer electrodes and probes (Trautmann et al., 2023). Our goal over the last three years has been to develop reliable methods to insert both probes, repeatedly, quickly, and acutely during each experimental session, to facilitate the study of 10s of neurons simultaneously in the awake NHP.

We describe two methods customized for rodent and NHP probes. For the rodent probe, we developed a method for insertion via a short guide tube eyelet, based on a concept developed by Matsuzaka et al. (2009). This method does not require a durotomy or probe sharpening. For the NHP probe, we have developed a recording chamber system with an artificial dura for repeated and easy insertion of a sharpened probe. Using these methods we have recorded activity from neurons in V2, V4 and prefrontal cortex (PFC) for several months in each case.

## 2. Materials and methods

### 2.1. Animal preparation

Three macaque monkeys, one female (Z: 6 kg) and two male (F: 9.2 kg, L: 13.4 kg), participated in these experiments. Animals were surgically implanted with custom-built head posts attached to the skull with orthopedic screws. After behavioral training, a metal ring was implanted on the skull surface of each monkey, followed in a subsequent surgery by a craniotomy and the installation of a plastic or metal recording chamber. Monkey Z had a chamber implanted over dorsal V4 and adjoining sulcal banks on the right hemisphere. Monkey F had chambers sequentially implanted over V2 on the left hemisphere and then V4 on the right hemisphere. Monkey L had two chambers simultaneously, one over dorsal V4 and the other over PFC for simultaneous study of the two brain regions. These chambers were first implanted over the left hemisphere and then over the right hemisphere, resulting in a total of 4 chambers. Chamber locations were determined based on stereotaxic coordinates and structural magnetic resonance images taken prior to headpost implantation. V2 and V4 recording sites were validated on the basis of the observed receptive field progression along the A-P and M-L axes of the chamber. In Monkey L, the chamber location could be confirmed after the durotomy by looking at the pattern of sulci inside the chamber. In Monkeys Z and F, penetration locations were visually confirmed after euthanasia.

For guide tube penetration and NHP probe penetration via native dura, it was critical to maintain a thin dural layer with periodic (once every 2-3 weeks) dural debridement under ketamine (with pain medication). All animal procedures conformed to National Institutes of Health guidelines and were approved by the Institutional Animal Care and Use Committee at the University of Washington (protocol # 4133-01).

### 2.2. Artificial dura chamber

In Monkey L we performed a durotomy roughly 4-6 weeks after the initial craniotomy when insertion of the NHP probe via native dura became difficult. The size of the durotomy varied across chambers (∼6-12 mm in diameter) but was at least 4 mm smaller in diameter than the craniotomy. We then placed a circular, silicone artificial dura (AD) to cover the exposed cortex. The AD was tucked under the native dura and glued to it around the edges with a thin layer of Vetbond (3M) applied using a sterile fine bristle paint brush (see Figure 1). The AD was ∼200 μm in thickness and was made by the Orsborn lab at the University of Washington. The AD was trimmed to a size that was ∼2-4 mm larger in diameter than the durotomy so as to have a generous overlap with the native dura and, where possible, tucked under the skull. Blood vessels and sulcal landmarks were visible through the AD for ∼1-2 months after durotomy, which facilitated electrode targeting. Periodically (∼ once every 2 weeks) as the glue crumbled and the AD became unglued, we reapplied Vetbond as needed until eventually the AD became untucked from the native dura and was unable to be reinserted. At this point we removed the AD and continued probe insertion via the tissue that had regrown under the AD. We were able to successfully insert the NHP probe for a duration of 3-4 months using this method.

**Figure 1.**
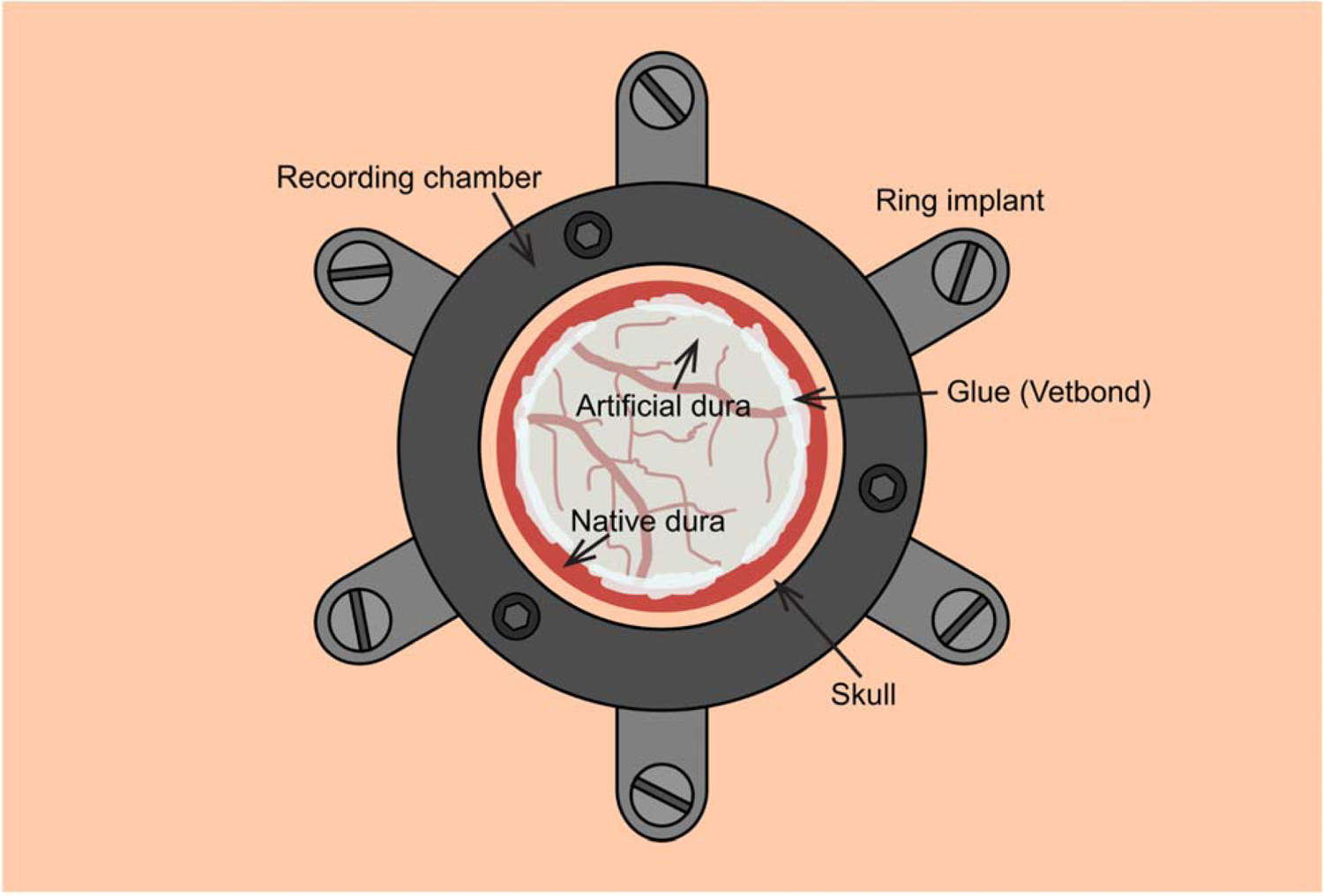
Top down view of the artificial dura (AD) chamber (not to scale). Our recording chambers have two parts: a low profile ring that is secured to the skull with titanium screws (pale gray) and a cylindrical chamber that is attached to the ring with screws (darker gray; see Figure 6 for side view). We first implant the ring and suture skin over it for healing. In a second surgery at least 6 weeks later, we perform a craniotomy and attach the recording chamber. The craniotomy is made as close to the edge of the chamber as possible using a piezo drill. Chamber inserts and caps are designed to provide a good seal In a subsequent surgery, we perform a durotomy, at least 4 mm smaller in diameter than the craniotomy. The AD is tucked underneath the native dura and glue (Vetbond, 3M) is applied around the edges with a fine-tipped brush. The surface of the brain with blood vessels and sulci can be seen underneath the transparent AD for approximately 1-2 months post-durotomy until tissue grows back underneath.

### 2.3. Chamber cleaning

For cleaning recording chambers with intact native dura, we typically flush the chamber first with 150-200 mls of 2% chlorhexidine diluted with sterile water (ratio of 1:3) followed by ∼50 mls of 0.9% saline solution at least 3 times per week. We used this procedure to maintain chambers in Monkeys Z and F where no durotomies were performed. In Monkey L, in order to discourage tissue growth on the dura before the durotomy, we used two strategies. At the end of each experiment, we flushed the chamber as described above and then applied a silicone gel (approximately 500µL in a 1:1 ratio of part A to part B; Dowsil 3-4680 Silicone Gel, Dow Corning) to serve as a dural sealant (Jackson and Muthuswamy, 2008). We also designed our chamber plugs and caps with o-rings to create a well-sealed chamber assembly.

For the AD chambers, beginning immediately after the durotomy and continuing until a thick layer of tissue had regrown over the exposed cortex, we flushed the chamber with ∼100mLs artificial cerebrospinal fluid (aCSF) (8.0661g NaCl, 0.208g KCl, 0.353g CaCl_2_*2H_2_O, and 0.549g MgCl_2_*6H_2_O per 1 L deionized water combined with 250mLs PBS) at least 3 times per week, and before and after every recording session. The aCSF was stored in a refrigerator and warmed in a water bath to body temperature (∼37 °C) before being used to flush the chamber.

### 2.4. Short guide tube construction

Guide tubes (Figure 2A) were constructed from 27 G hypodermic needles (BD or EXEL) and epoxy glue. Hypodermic needles were cut to a length of 3-4.5mm from the taper tip using an electric hand dremel and deburred using an 18 G or 23 G needle. Then a small blob of epoxy glue was placed close to the blunt end of the guide tube to serve as an anchor against the dura. Roughening the surface of the guide tube prior to epoxy placement provided a better bond. Care was taken to ensure that a 1-2.0 mm length of guide tube from the sharp tip was left uncovered by glue. Care should also be taken to ensure that epoxy does not flow into the guide tube.

**Figure 2.**
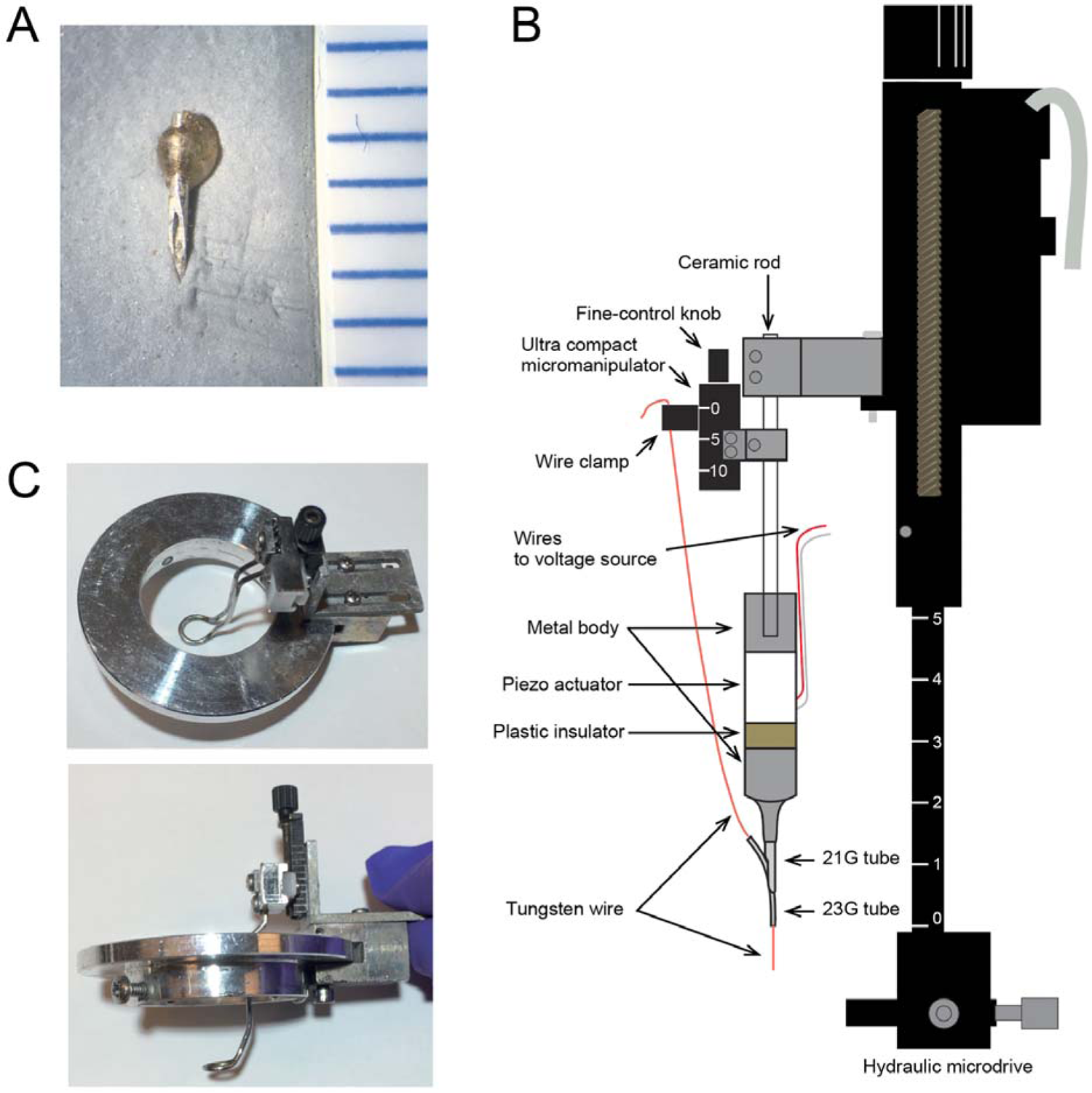
Guide tube insertion parts. ***A.*** An example of a short guide tube used in recording. The scale shown on the right is in millimeters. A 3.5 mm guide tube is shown with an epoxy blob (distal to the sharp tip). A 2 mm length of guide tube at the tip is clear of epoxy. ***B.*** Guide tube launcher assembly secured to the hydraulic microdrive (black). See Methods for details. ***C.*** Side and top view of the presser foot mounted on the chamber lid (silver disc) using an ultracompact micromanipulator (black).

### 2.5. Guide tube launcher

Schematic drawing of the guide tube launcher assembly is shown in Figure 2B. The device consists of 23 G and 21G hypodermic needles (BD), a piezoelectric actuator, an ultracompact micromanipulator (Narishige, MO-903B), custom-made metal body and a ceramic rod for securing to the hydraulic microdrive (Narishige, MO-97A). The piezoelectric actuator (KEMET) provided axial vibration to facilitate cutting soft tissue as we inserted the guide tube (Noda et al., 2011; Barnett et al., 2016). We used a tungsten wire (200μm tungsten electrode, FHC) to hold the short guide tube and the wire tip was shaved so as to provide a snug fit to the inner diameter of the 27G needle. The far end of the wire was connected to an ultracompact micromanipulator to facilitate retraction of the wire and release of the short guide tube.

### 2.6. Guide tube insertion procedure

Each day we identified our target recording location by placing a custom-built plastic grid with 1 mm spacing in the recording chamber, inserting a needle dipped in India ink through the desired grid location and marking the underlying dural location. The grid was then removed and a custom-built metal presser foot was placed against the dura mater so as to gently stabilize the dura centered at the marked location (Figure 3A). The wire presser foot was held with an ultracompact micromanipulator secured to the chamber (Figure 2C). This allowed daily adjustment of the presser ring depth to apply just enough pressure on the dura. Then, the guide tube launcher assembly was advanced toward the dura using a hydraulic microdrive until the short guide tube pierced the dura (Figure 3B). Once the short guide tube was anchored against the dura with the epoxy glue blob, the tungsten wire inside the guide tube was slowly retracted using the ultracompact micromanipulator (Figure 3C). This detached the short guide tube from the guide tube launcher which was then retracted using the hydraulic microdrive (Figure 3D). If the guide tube had pierced through the dura, clear cerebrospinal fluid (CSF) could be observed coming up from the guide tube. We activated the piezo drive for axial vibration during guide tube placement as needed. The piezo drive is wired to an amplifier (PD200 60W Voltage Amplifier, PiezoDrive) and signal generator (15MHz High Precision DDS Signal Generator, Koolertron). The frequency of the signal generator was set to below 500 Hz and the output level to 0-5V. The output voltage of the amplifier was less than 115V.

**Figure 3.**
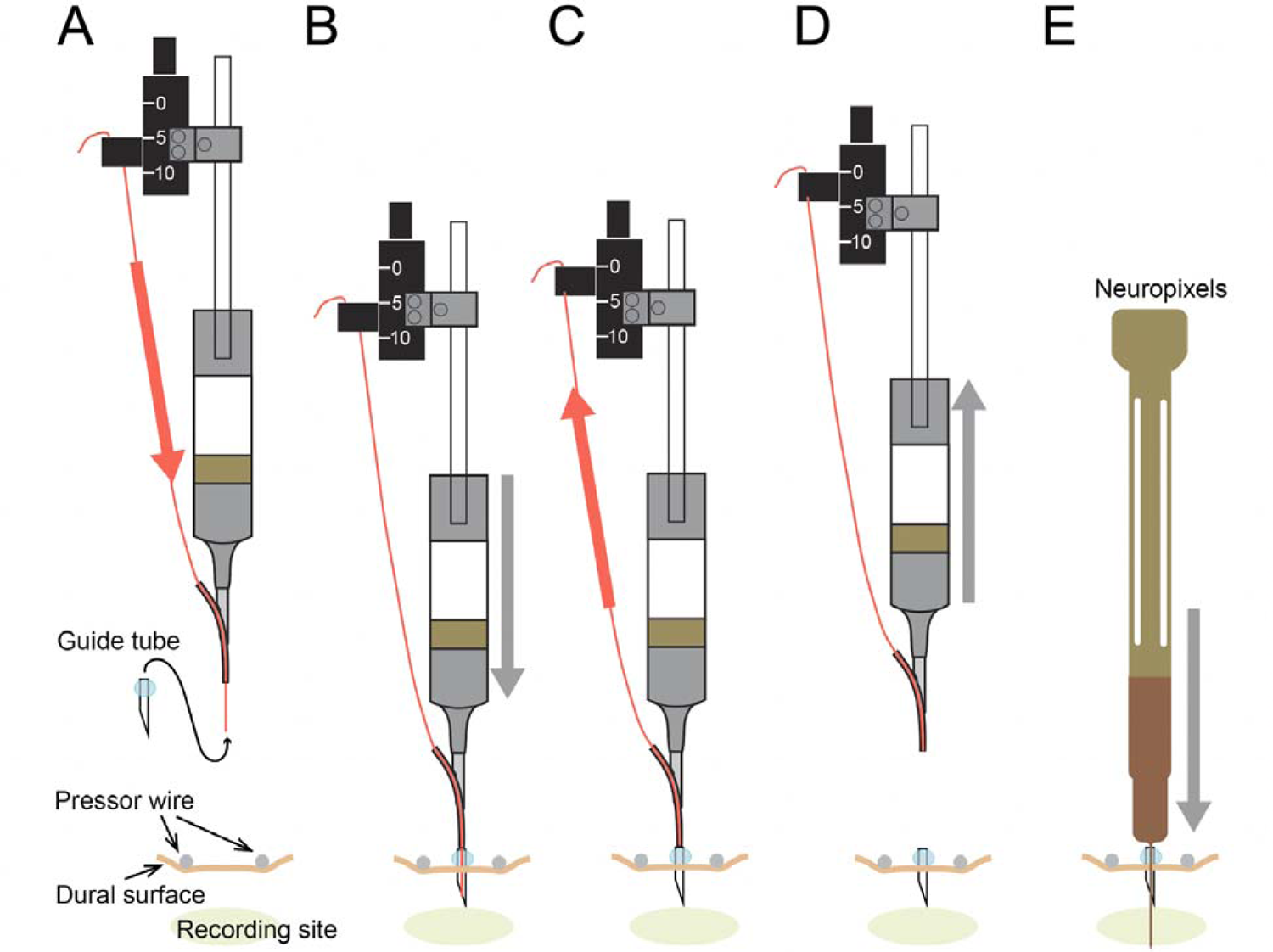
Schematic of guide tube launcher set up and insertion procedure. ***A.*** Preparation of the guide­ tube launcher. The tungsten wire (red) was advanced out of the 23G tubing using the ultracompact micromanipulator and tweezers. Then, the short guide tube was mounted securely onto the tungsten wire with tweezers. The guide tube and tip of the guide-tube launcher were sterilized with Cidex OPA and rinsed with saline before insertion. When ready to insert, the presser foot was positioned against the dura encompassing the desired recording site (green oval). ***B.*** Inserting the guide tube. The entire guide tube launcher assembly was lowered using the hydraulic microdrive until the short guide tube pierced the dura and the epoxy blob was anchored against the dura. *C.* Releasing the guide tube from the tungsten wire. Using the ultracompact micromanipulator, the tungsten wire was retracted out of the short guide tube. ***D.*** Retracting the guide tube launcher. The guide tube launcher assembly was retracted using the hydraulic: microdrive leaving the guide tube in the native dura. The guide tube launcher assembly was then removed from the hydraulic microdrive and replaced with a dovetail holder with the Neuropixels probe. *E.* Inserting the probe. Data acquisition cables were plugged into the probe headstage and the reference and ground were wired to the headpost. Then, the probe was finally inserted in the recording site through the dural eyelet. See Methods for more details.

### 2.7. Rodent probe insertion

The rodent probe was mounted on a dovetail holder (Sensapex) customized to attach to the hydraulic microdrive. The tip of the probe and guide tube hole were aligned under magnification (telescopic magnifier system, Zeiss K 4.5x/350mm). Data acquisition cables were plugged into the headstage and the reference and ground were wired to the headpost prior to probe insertion into the guide tube. The probe was inserted into the cortex through the short guide tube using the hydraulic microdrive (Figure 3E).

During insertion, the activity map on spikeGLX was continuously monitored by the experimenter and the activity progression along the length of the probe guided decisions on depth of insertion. We advanced the probe until robust activity was evident in the middle of bank 0 (the bottom one-third of the probe) or the top of the guide tube was located at the middle of bank 2 (top one-third of the probe).

### 2.8. Sharpening the NHP probe

To sharpen the NHP probe we used a modified version of a rodent probe sharpening protocol originally developed by the Steinmetz lab at the University of Washington (https://github.com/cortex-lab/neuropixels/wiki/Sharpening). Before sharpening each probe, we ran diagnostic tests on the probe to ensure normal functioning. This was done by connecting the probe to the recording system and starting an acquisition and/or running BIST tests. Then, the probe was mounted on a 3D printed holder (light gray in Figure 4) designed to mate with the holder on the micro-grinder (Narishige EG-45). The probe was positioned with contacts facing up (i.e. the silver surface on the probe base is facing up) and the flat surface of the probe as parallel to the grinding surface as possible (Figure 4A). Since the grinder spins in one direction, it is important to position the probe such that the grinding wheel approaches the probe from shaft to tip (see Figure 4B) rather than in the opposite direction. With the probe tip well above and oriented parallel to the grinding surface, it was gradually lowered to an angle of 25°. The grinding wheel was then started at the lowest speed (10-20 on the 0-100 range of the dial; ∼ 200 rpm), and the probe was carefully lowered (using the coarse and then fine control) until it just made contact with the grinding wheel. During this entire process we monitored the probe tip under magnification (stereo microscope,

**Figure 4.**
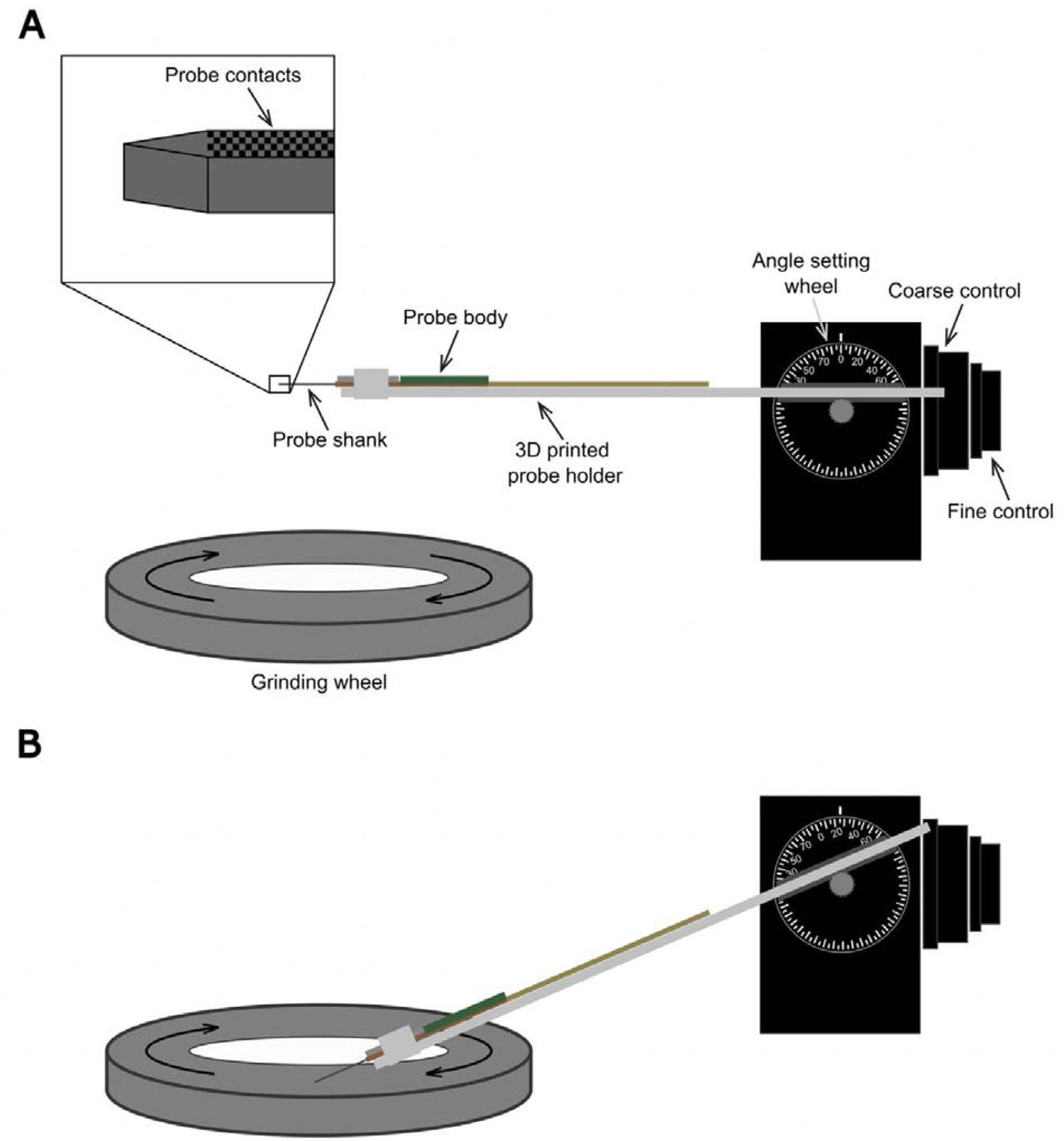
NHP probe sharpening set up. ***A.*** An NHP probe positioned in the Narashige grinder holder pre-sharpening. Probe contacts should be facing up during sharpening and the probe should be made as level as possible during this stage. ***B.*** An NHP probe during sharpening. The angle is set to 25” and the probe is lowered onto the surface of the grinder while the disk is slowly spinning before ramping up the speed to 50. The probe should be positioned on the grinding wheel so that the grinding wheel approaches the probe from shaft to tip as seen here.

Leica M80). It is advised to position a light source such that the probe’s shadow on the wheel can be observed meeting the tip of the probe. Once the probe made contact with the grinding surface (i.e. when the shadow and the tip of the probe were touching), it was lowered one additional turn (= ∼⅓ mm) on the fine control wheel of the grinder. Then, the speed was gradually increased to 50 on the dial (= ∼ 800-900 rpm) while monitoring the probe under the microscope for any up/down bouncing, which is a sign that the speed is too high. If bouncing was observed, since this can damage the probe, we raised the probe away from the grinding surface, lowered the speed back down and repeated the approach as described above. At a speed of 50, the grinding process continued for 5-6 minutes, at which point the probe was slowly raised off the grinder while it was still spinning. We then stop the grinder, raise the probe further, and reset the holder angle back to 0° to inspect the tip under magnification and determine if additional sharpening is needed (see Figure 5), repeating the sharpening process for an additional 1-2 minutes at a time if necessary. A total sharpening duration of 7-8 minutes has worked well with many probes, facilitating easier penetration of the native dura or AD. Once sharpening was complete the probe was retested to make sure it was not damaged during sharpening.

**Figure 5.**
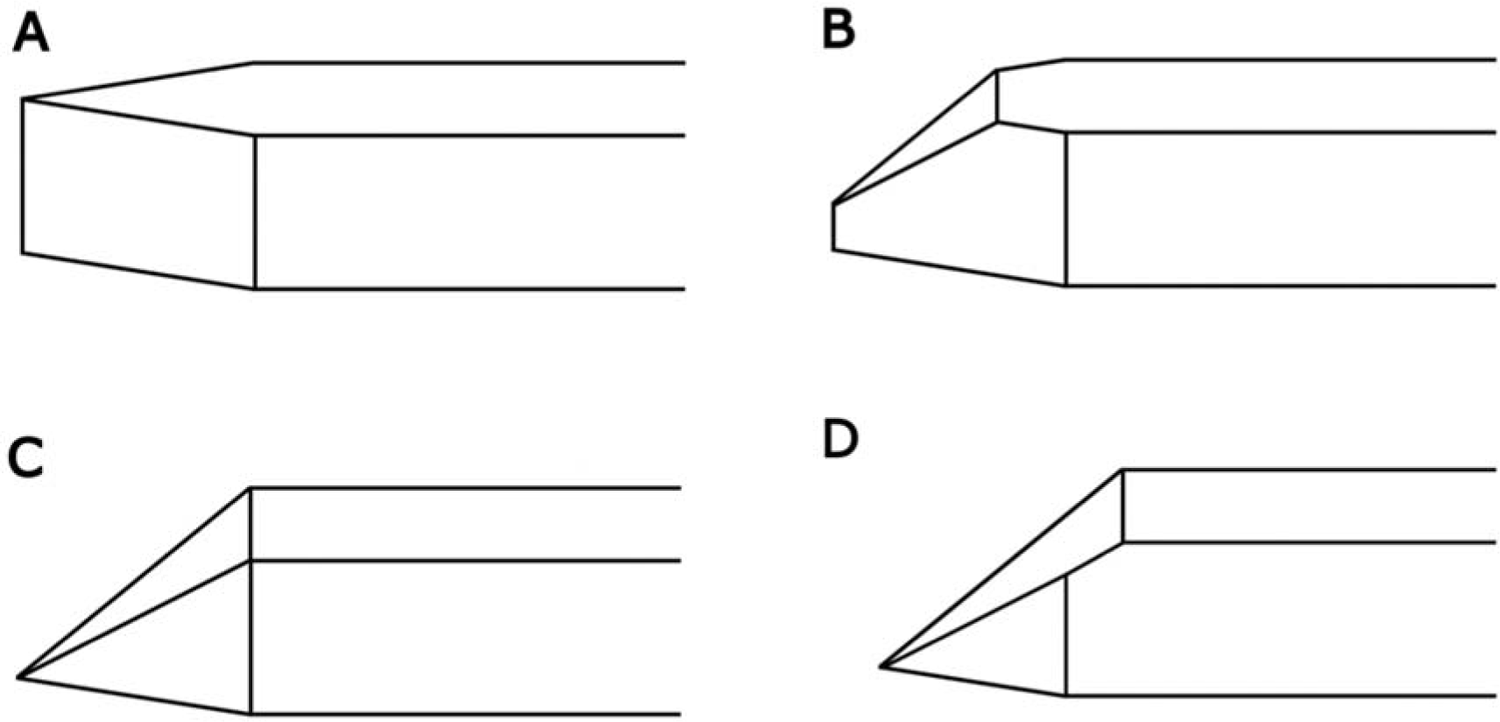
NHP probe at various stages of sharpening. ***A.*** Unsharpened NHP probe. The electrode contacts are located on the bottom (not visible) surface of the probe (note that these diagrams are flipped from how the probe would actually be positioned during sharpening, where probe contacts would be positioned upwards). ***B.*** Partially sharpened probe. This probe is not as sharp as it could be and further sharpening should be performed. ***C.*** Very sharp probe. Sharpening is complete at this stage, when all surfaces narrow to a single point. ***D.*** Overly sharpened probe. Sharpening was likely overdone and the probe might be broken at this stage.

**Figure 6.**
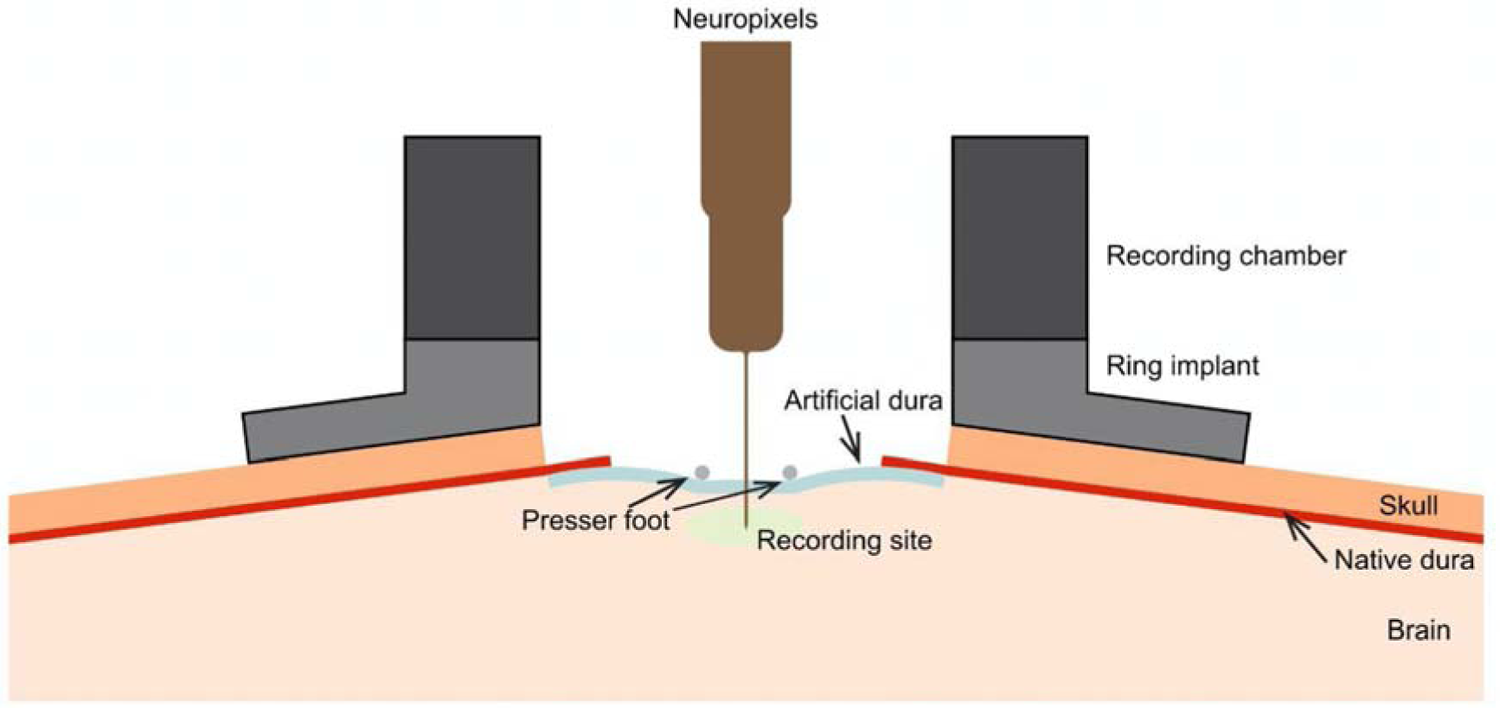
Side view of the AD chamber during probe insertion. The presser foot is lowered to apply gentle pressure on the AD and underlying cortex encompassing the desired recording site. In tall and narrow chambers, visualizing bending in the probe can be difficult due to limited lines of sight.

### 2.9. NHP probe insertion

We use the same process to insert NHP probes through native dura, AD and regrown tissue. Below, we describe the insertion process through an AD, but the terms AD, native dura, and regrown tissue can be used interchangeably in this section, as the insertion method is largely the same. NHP probes were mounted on a dovetail holder (Sensapex) customized to fit the hydraulic microdrive. To ensure stability of cortex during recording and assist with probe insertion, a metal presser foot (Figure 2C) was lowered onto the surface of the AD using the ultracompact micromanipulator until gentle pressure was applied on the surface but did not appear to cause any discomfort to the animal. Too much pressure with the presser foot could result in poor spike yield in the recording, with the activity of the neurons along the probe looking ‘silent’. Landmarks within the chamber such as sulci and blood vessels, the durotomy border, and other tissue formations were used to guide probe targeting across days. Data acquisition cables were plugged into the headstage and the reference and ground were wired to the headpost prior to probe insertion. The probe was lowered into the chamber with the coarse drive on the hydraulic microdrive until it neared the surface of the AD, after which the fine drive was used and the x/y recording location was targeted under magnification. The probe was inserted through the AD and into the cortex using the fine control on the drive. The typical speed of insertion used was ∼10µm/second. We have used both a manually controlled and a motorized hydraulic microdrive for insertion and both work equally well; the choice is largely up to experimenter preference and experience. During insertion the activity map on spikeGLX was continuously monitored by the experimenter and the activity progression along the length of the probe guided decisions on depth of insertion. We advanced the probe until robust activity was evident in the middle of bank 0 (the bottom 1/3rd of the probe). During insertion the probe shank was also carefully and continuously observed under magnification to monitor for any bending of the shank, which can lead to breakage if the probe continues to bend while being advanced. With a fresh craniotomy or recent durotomy/AD placement, probe bending was not much of an issue, but as tissue thickened on the dura or underneath the AD, bending became a significant problem and probe insertion was more difficult. The probe bends primarily along the long axis which can be difficult to detect depending on the geometry of the recording setup and angles of visibility. A successful method for detecting bending was to watch for any slight lateral movements of the probe shank as it often appeared to wiggle with slight movements in the dura or tissue when beginning to bend. With both the dural eyelet and AD chamber methods, rather than waiting after probe insertion, we found that stable recordings could be reliably achieved if the probe is inserted deeper than necessary and then retracted until there is a visible change in the activity map on spikeGLX. This retraction also releases any stress/bending of the probe which on some occasions can be visualized under magnification.

All details about the visual display, data collection hardware and offline analyses are described elsewhere (Bigelow et al., 2023; Namima et al., manuscript submitted for publication).

## 3. Results

We have developed multiple methods for the acute insertion of Neuropixels probes into the awake monkey cortex. Our first strategy was designed for the rodent probe based on Matsusaka et al. (2009). Subsequently we developed methods for the NHP probe both with native dura and the use of an artificial dura chamber. Below we describe the time needed for probe insertion, failure modes and broken probe retrieval.

### 3.1. Rodent probe experiments

We performed acute recording from the cerebral cortex using a rodent probe in two awake macaque monkeys: 85 penetrations in Monkey Z over a 1 year period targeting area V4 on the right hemisphere and 52 penetrations in Monkey F over 5 months in a chamber targeting area V2 on the left hemisphere. In both cases, we stopped recording when we had collected sufficient data and not because recording quality had deteriorated. Data quality and yield were good even on the last day of the recordings. For some results from these penetrations please see Bigelow et al., 2023.

#### 3.1.1. Penetration of dura

With our dural eyelet method, the time needed for insertion of the guide tube through dura varied across days. Under ideal conditions of visibility, thinness of dura and sharpness of the guide tube tip, this was a fast process that required 10 minutes. Care should be taken to pierce all layers of the dura. An indication of this could be the emergence of cerebrospinal fluid (CSF) through the dural eyelet. When the dura was thick and the guide tube simply tented the dura rather than pierce it, we used piezo axial vibration to facilitate piercing without excessive force. We prepared several guide tubes with different lengths prior to recording and selected a guide tube with the appropriate length. For sessions without prior knowledge about dural thickness, we started with a shorter guide tube and increased guide tube length if piercing the dura failed with the procedures described above. When we had made a penetration in a nearby cortical location in the preceding few days, we used the same guide tube or another of similar length.

#### 3.1.2. Failure of probe insertion

After creating a dural eyelet by placing the short guide tube, the rodent probe was inserted through the eyelet under magnification. It is critical at this point to proceed carefully and watch for bending or buckling. On some occasions we failed to advance the probe into the cortex, for a few reasons. First, it is critical to use a holder that is designed to hold the probe parallel to the axis of motion. The first iteration of our probe holder was flawed in this regard, which resulted in many broken probes. Second, the orientation of the short guide tube may be at an angle relative to the axis of the probe, causing the probe tip to hit the wall of the guide tube and buckle or bend when advanced. Third, CSF which comes up the guide tube after the dura has been successfully pierced may dry, clogging the eyelet. This most commonly occurs when the guide tube is of optimal length and the hole on the needle bevel lies just above the surface of cortex. In this case, inserting a probe through the same length guide tube more quickly on the immediate next attempt could solve the issue. Care also needs to be taken to ensure that the 10 mm probe is not fully inserted through the guide tube causing the base to touch the top of the guide tube (See Rodent probe insertion in Materials and Methods).

#### 3.1.3. Probe contacts available for recording

With this method, since the guide tube covers some of the probe shank, it is important to consider the length of the probe available for recording. Our guide tubes were typically 3-4 mm long, so the available probe length for insertion into the cortex was ∼5-6 mm after allowing for 0.5-1 mm of probe above the top of the guide tube. If the probe contacts face the needle bevel, this further exposes another 1mm of the probe.

### 3.2. NHP probe experiments

In Monkey F, we performed 70 and 6 penetrations through native dura into V4 of the right hemisphere and V2 of the left hemisphere, respectively. Immediately after the craniotomy, the NHP probe easily pierced native dura. With time, as the dura thickened, we performed periodic topical cleaning of the dural surface under sedation, which facilitated the insertion of the probe. In Monkey L, we performed 59 sessions of acute recordings from two probes simultaneously, one in the PFC and another in area V4, as well 19 sessions in which only one of the two areas was recorded from. In these experiments, immediately after craniotomy, we inserted the NHP probe through native dura. As the native dura thickened, we performed a durotomy and placed an AD in each chamber. Of the dual recordings, 5 sessions were through native dura and 54 were through the AD or regrown tissue, while 14 of the single probe recordings were through native dura and 5 were through the AD or regrown tissue. These procedures were replicated in two pairs of chambers in the left and right hemispheres of the same animal (Monkey L).

#### 3.2.1. Probe insertion

The time needed for NHP probe insertion into the cortex varied across days. In a fresh craniotomy or with a recently placed AD, probe insertion proceeded with relative ease. Under these conditions, there was minimal probe bending and insertion could take as little as 10-15 minutes. As the dura toughens or tissue grows under the AD, probe bending is a significant problem which can prolong the insertion process. We have found that a few strategies can facilitate probe insertion through thicker native dura or regrown tissue (both with an AD and after the AD had come unglued and been removed from the chamber). One method that has been successful when tissue has begun to grow but has not yet become extremely thick and tough was to advance and retract the probe in a sewing machine type motion, slowly going up and down about 200-500µm at a time, while maintaining an insertion speed of ∼10µm/sec and carefully observing the probe for bending – stopping and retracting again when bending is observed. Frequently with this method we are able to penetrate farther with each advancement before bending occurs again and eventually observe the probe ‘punch through’ the remaining tissue/dura and stop bending, allowing us to continue insertion normally. When the tissue has thickened and the sewing machine method fails, we have successfully inserted the probe by lowering the probe until a slight bend is observed and then waiting until the probe pops through on its own. This method, however, can be time consuming and has resulted in probe breakage on two occasions. On both occasions, a forceful animal movement preceded the breakage. If neither of the above techniques are successful, sometimes retracting the probe and attempting to insert in a nearby location can be effective, likely due to natural heterogeneity in tissue structure or thickness throughout the chamber.

#### 3.2.2. Failure of probe insertion

Insertion of NHP probes can fail for a variety of reasons, either by termination of the insertion attempt due to too much elapsed time or by breakage of the probe shank. Insertion through tough or thick native dura or regrown tissue can be very tedious and time consuming which, depending on the temperament of the animal can cause them to become impatient and shake their body inside the primate chair or to not work later on in the recording session, making long insertion periods not ideal. Additionally, as described in the above section, a combination of a large bend in the probe and the animal shaking or moving forcefully in the chair can result in the probe breaking. This is the primary instance in which we have encountered probe breakage while using this method.

#### 3.2.3. Probe contacts available for recording

The AD chamber method avoids the problem of covering probe contacts with the guide tube so the entire length of the probe is available for recording. However, since a neo-membrane can grow on top of the cortex in varying thicknesses and there can be fluid trapped in between the AD and the surface of the underlying tissue, it is possible that the first sign of cells will appear 5-6 mm below the surface of the AD. We have never had an issue with not being able to reach active cortex before inserting the entire 10 mm shank of the probe but this could theoretically be a problem with this method.

### 3.3. Broken probe retrieval

The rodent and NHP probes can break for several reasons. Continuing to advance a bending probe could lead to breakage. Inadvertent taps orthogonal to the long axis can also break the probe. On a few occasions, our probes have broken while partially in cortex. In this case, the probe tip was partially left in the cortex and/or the guide tube and we followed the procedure described below to remove the broken probe tip. We pulled up the probe holder using the hydraulic microdrive and replaced the broken probe and holder with a Q-tip. We suctioned out the fluid in the recording chamber with a vacuum or absorbed it with Q-tips and allowed the exposed part of the probe tip to dry. Next, we applied a drop of Kwik-sil (World Precision Instruments) to the Q-tip attached to the microdrive and gently (with the fine adjustment) extended it until the Kwik-sil ball was in contact with the probe piece in cortex. We waited 5 minutes and gently retracted the Q-tip with fine adjustment of the microdrive. Repeat the process as many times as necessary if the first attempt is not successful but note that you do not want to touch the broken probe or push the probe further into the cortex while attempting to remove it. This procedure has worked well to remove broken both rodent and NHP probes.

### 3.4. Failure of probe retraction

With the AD chamber, we have encountered difficulties with retracting the probe on a few occasions primarily due to drying of the chamber. This can cause material from underneath the AD to get stuck to the probe, tenting the AD, which has on one occasion resulted in a hole being torn in the AD. To prevent this, we drip artificial cerebrospinal fluid (aCSF) into the chamber at the end of the recording immediately before probe retraction using a small syringe and needle (we use 21G needles and a 10ml syringe) until the recording site is completely submerged. With aCSF in the chamber we have not experienced any issues with the AD tenting during probe retraction. With the short guide tube, we had a few occasions where the guide tube came up with the probe when the probe was retracted. This was because fluid in the guide tube dried during the experiment causing the guide tube to adhere to the probe shank. Soaking that retracted probe in saline or Tergazyme solution (1% concentration, Alconox) for several hours can release the probe from the guide tube. In our case, this did not cause any damage to the probe.

### 3.5. Data quality

Figure 7 compares results from various example recording sessions using the rodent (A-C) and NHP probes (D-F). With both probes we typically waited at least 5 minutes, and often ∼30 minutes, prior to the start of our main experiments to allow for stability of the recording preparation. Qualitative comparison from our recording sessions suggest similar yield and stability with both probes. More detailed analyses are needed to rigorously compare these methods.

**Figure 7.**
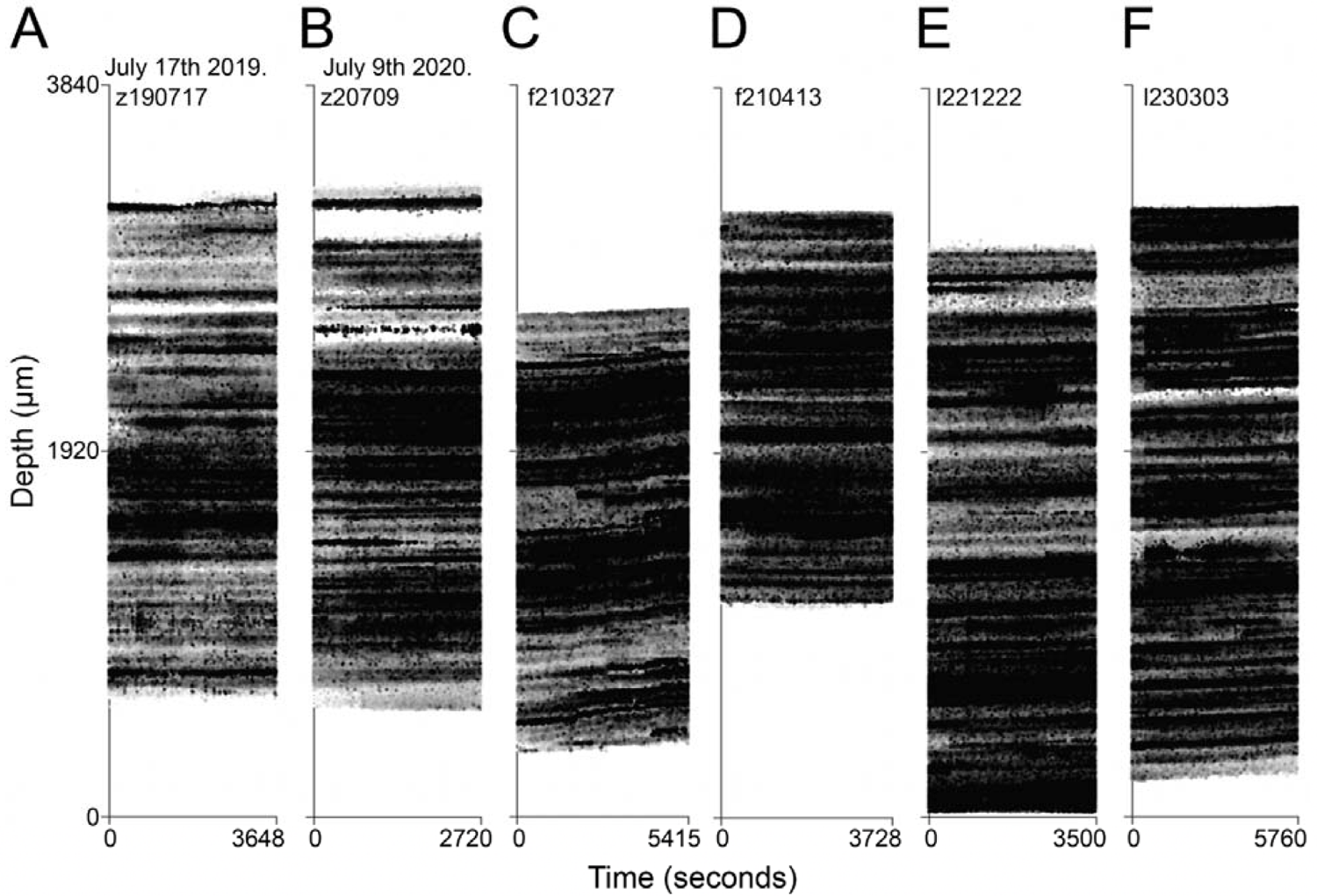
Cortical activity band obtained with rodent (A-CJ or NHP probes (D-F). Individual spikes (grayscale dots) are plotted as a function ofrecording time (X axis) along the probe depth (Y axis). The dot grayscale indicates relative amplitude of spike waveforms within a session, with darker dots indicating larger amplitudes. *A.* Driftmap for a recording session in V4 with a rodent probe performed for over one hour. Y = 0 corresponds to the deepest recording contact in bank 0. ***B.*** Driftmap for a recording session in V4 cortex in the same recording chamber as in A (from Monkey Z), also with a rodent probe. Recording B was performed roughly one year after recording A (see dates) but the data yield was similar. ***C.*** Another example from a rodent probe recording in V2 with a short guide tube from Monkey F. Probe stabilization was not sufficient so spike waveforms can be seen drifting up away from the probe tip during the entire −90 minute recording session, but the drift of spikes was small enough for Kilosort to successfully spike sort. *D.* Driftmap from a recording session in which an NHP probe was inserted into area V4 through native dura. Upon insertion, the probe was slightly retracted to gain better stabilization. This is an example of a recording session with minimal drift for the entire hour-long recording duration. *E.* Driftmap from a recording session with an NHP probe inserted in V4 of Monkey L through native dura approximately 2 weeks post-craniotomy. *F.* NHP probe recording from V4 of Monkey L through artificial dura approx. 2 months after the durotomy and AD placement surgery. In all panels, recording data for different experiments were concatenated post hoc. As a result, vertical banding patterns may be visible on the activity map, reflecting discontinuities of units across time. Driftmaps in A and B-F were created by applying ksDriftmap.m script (cortex-lab) to outputs ofK.ilosort2.5 and Kilosort 2.0, respectively.

## 4. Discussion

Our goal was to develop methods to insert rodent and NHP Neuropixels probes into the awake monkey cortex. Using the dural eyelet method for rodent probes and an AD chamber for NHP probes we have successfully collected data in nearly 300 recording sessions across 7 chambers in 3 monkeys. Below we discuss pros and cons of each method and our suggestions for improvements.

### 4.1. Relation to prior studies

Our methods complement a few previously published reports that have used Neuropixels probes in the macaque monkey. In the first published report, Trautmann et al., (2019) achieved success by pre-puncturing the dura with a needle followed by insertion of the rodent probe. In the anesthetized animal, Trepka et al., (2022) inserted rodent probes after reflecting the dura to expose cortex. Bauer et al., (2023) used a three stage method – pre-puncture of dura, guide-tube insertion and probe insertion – with both rodent and NHP probes. Others (e.g. Wang et al., 2021) have attempted to reinforce the probe shank to facilitate robust insertion.

Similar to Trautmann and colleagues, our first strategy with rodent probes was pre-puncture. However, we had a high failure rate with probe insertion, either because the probe tip failed to target the punctured location even though this was done under magnification or because the puncture location closed before probe insertion. We believe that our dural eyelet method improves upon the pre-puncture strategy because the eyelet can be easily targeted and the probe may be inserted with less friction. More broadly, the advantages of this method are that it works with an intact dura prep (no durotomy is required), and can work with extremely fragile, short probes. In terms of downsides, the probe insertion itself can sometimes take a long time and the guide tube may cause cortical damage or shield contacts and reduce yield from the superficial cortex. Overall, this dural eyelet method could make it possible for NHP researchers to use any short, fine high density probes developed for use in rodents, in the monkey model. This could accelerate data collection in the monkey.

With regards to the AD chamber approach, advantages include fast and easy probe insertion and minimal damage to the superficial cortex compared to pre-puncture and the dural eyelet method. In terms of downsides, removal of native dura can raise the risk of infection and require greater vigilance. Also, the NHP probe may be unable to pierce through blood vessels or other thickened tissue and may become stuck and lead to breakage.

### 4.2. Troubleshooting and future modifications

Once the probe is securely in the cortex, the activity on the probe may be weak if (i) the presser foot is applying too much pressure or (ii) the probe has repeatedly been targeting the same piece of tissue across many days. In these cases, (i) retracting the presser foot to release some pressure or (ii) targeting an adjacent piece of tissue could fix the issue.

Future studies could improve upon our procedures to address some key concerns. One concern is the anchor provided by the epoxy glue blob. While the epoxy is easy to apply and it works well, it is possible that it could lose its bond to the guide tube when it is pushed up against the dura causing the guide tube to be forced deeper into the cortex. A more robust anchor could be achieved by silver soldering.

Asymmetries in the epoxy (or solder) could lead to the guide tube anchoring at an angle to the dura which can cause probe breakage. Future attempts could create a better anchoring system for the short guide tube. A second concern is the damage to the superficial cortex caused by the guide tube. Knowing the best guide tube length could be easier if a tungsten electrode is positioned inside the short guide tube during insertion to detect the first sign of neural activity. This strategy could help to identify an appropriate guide tube length that would not invade the superficial cortex. In addition to avoiding damage this would also be important for more accurate estimates of neuron depth in the cortex.

Finally, given that the NHP probe can pierce native dura immediately after a craniotomy, one effective strategy could be to develop methods that could retard tissue growth and thickening of the NHP dura to facilitate probe insertion without the need for pre-puncture, a dural eyelet or an AD chamber.

## Acknowledgements

We thank Amber Fyall for animal care and behavioral training, WaNPRC Instrumentation Services for hardware support, and Tony Bigelow and Sofia Beaufrand for help with experiments.

## CRediT authorship contribution statement

Tomoyuki Namima: Conceptualization, Methodology, Investigation, Visualization, Writing – original draft, Funding acquisition. Erin Kempkes: Methodology, Investigation, Visualization, Writing – original draft. Bob Smith: Resources. Anitha Pasupathy: Supervision, Conceptualization, Methodology, Investigation, Visualization, Writing – original draft, Funding acquisition.

